# First Eurasian cases of SARS-CoV-2 seropositivity in a free-ranging urban population of wild fallow deer

**DOI:** 10.1101/2023.07.07.547941

**Authors:** Kevin Purves, Hannah Brown, Ruth Haverty, Andrew Ryan, Laura L. Griffin, Janet McCormack, Sophie O’Reilly, Patrick W. Mallon, Virginie Gautier, Joseph P. Cassidy, Aurelie Fabre, Michael J. Carr, Gabriel Gonzalez, Simone Ciuti, Nicola F. Fletcher

## Abstract

Severe acute respiratory syndrome coronavirus-2 (SARS-CoV-2) infects wildlife. Recent studies highlighted that variants of concern (VOC) may expand into novel animal reservoirs with the potential for reverse zoonosis. North American white-tailed deer are the only deer species in which SARS-CoV-2 has been documented, raising the question whether further reservoir species exist as new VOC emerge. Here, we report the first cases of deer SARS-CoV-2 seropositivity in Eurasia, in a city population of fallow deer in Dublin, Ireland. Deer were seronegative in 2020 (circulating variant in humans: Alpha), one animal was seropositive in 2021 (Delta variant), and 57% of animals tested in 2022 were seropositive (Omicron variant). Ex vivo, a clinical isolate of Omicron BA.1 infected fallow deer precision cut lung slice type-2 pneumocytes, also a major target of infection in human lungs. Our findings suggest a change in host tropism as new variants emerged in the human reservoir, highlighting the importance of continued wildlife disease monitoring and limiting human-wildlife contacts.

**Teaser:** This study is the first report of SARS-CoV-2 seropositivity in fallow deer, highlighting expansion of viral variants into new host reservoirs.

## 1. Introduction

Severe acute respiratory virus 2 (SARS-CoV-2) is a novel *Coronaviridae* family member (genus *Betacoronavirus*, subgenus *Sarbecovirus*) which possesses an enveloped viral particle and a positive-sense, single-stranded RNA genome of ∼30 kb (*1*). This pathogen is responsible for the COVID-19 pandemic which has led to approximately 7 million deaths in laboratory-confirmed cases (*2*) globally, and replicates in multiple mammalian species in addition to humans (*3, 4*). As well as zoonotic spread, anthroponosis (reverse zoonosis) is a concern with spillback of virus from humans-to-animals representing a potential route of virus spread and novel variant emergence (*5*). Recent reports of SARS-CoV-2 infection of white-tailed deer (*Odocoileus virginianus*) in the United States and Canada (*6-13*) have revealed high seroprevalence rates in free-ranging white-tailed deer. Moreover, viral shedding from infected animals and deer-to-deer transmission has been reported, with multiple lineages similar to contemporaneously-circulating human SARS-CoV-2 genomes detected within infected deer, indicating potential reverse zoonoses (*10, 12, 13*). This has raised concerns that other wild deer species may be susceptible to infection with SARS-CoV-2, and the World Organisation for Animal Health has recommended monitoring of deer species to better understand whether there is a likelihood of these species representing a reservoir for SARS-CoV-2 infection and potential for spread to other mammalian species, including humans (*14*).

To date, there is limited information on the susceptibility of European deer species to SARS-CoV-2 infection. Initial surveillance reports in the UK, Austria and Germany have not identified evidence of exposure to SARS-CoV-2 in European deer species, including fallow deer (*Dama dama*), red deer (*Cervus elaphus*), roe deer (*Capreolus capreolus*), sika deer (*Cervus nippon*) and other species, although surveillance has not been reported beyond 2021 (*15, 16*). Whether infection of these species does not occur, or is extremely rare, is unclear. Interactions of these wild, free-ranging species with humans may be low and therefore there may be limited opportunities for transmission events (*15*). However, a recent study reported expression of the viral primary receptor, angiotensin-converting enzyme-2 (ACE2), in bronchiolar epithelium of several deer species, including fallow deer, roe deer, red deer and muntjac (*Muntiacus reevesi*) (*17*). This raises the possibility that further deer species may be susceptible to SARS-CoV-2 infection, particularly with the emergence of novel, highly-transmissible variants of concern (VOC) in humans which may have expanded host tropism compared to ancestral viruses (*18*).

We have conducted a viral surveillance study in a wild free-ranging population of fallow deer in the largest urban park in Europe, the Phoenix Park in Dublin, Ireland. The Park receives up to 10 million visitors per year, and in the last decade tourists have increased their contact with the deer (*19, 20*). Park visitors have repeatedly been documented hand-feeding the deer population, leading to numerous direct interactions, e.g. people touching the deer while feeding them (*19*). The entire population of deer are constantly monitored via behavioural observations, with a precise estimate of the human-deer contact rates at the individual animal level and which are assigned a begging rank ranging from deer avoiding human contact to those consistently begging for food (*19*).

We investigated whether this peri-urban fallow deer population had evidence of SARS-CoV-2 infection during the period 2020-2022. A comprehensive set of tissue samples (retropharyngeal lymph nodes, tonsils, nasopharyngeal mucosa, caecal content) and sera were collected from animals culled as part of population control measures. Genome detection by qRT-PCR targeting the SARS-CoV-2 envelope (E) gene, as well as neutralising antibody titres using quantitative SARS-CoV-2 virus neutralisation test (sVNT), pseudovirus and infectious virus neutralisation assays were performed. SARS-CoV-2 serostatus was then correlated with the begging rank of each animal. Finally, the ability of SARS-CoV-2 isolates representative of the ancestral and Omicron variants to infect fallow deer precision cut lung slices and tracheal explants was evaluated.

## 2. Methods

### 2.1 Experimental design

This study aimed to investigate whether a peri-urban fallow deer population based in a European capital city, with defined interactions with humans, had evidence of SARS-CoV-2 infection during the period 2020-2022. Following culling for population control purposes, retropharyngeal lymph nodes, tonsils, nasopharyngeal mucosa, caecal content and sera were collected. Genome detection by qRT-PCR targeting the SARS-CoV-2 envelope (E) gene was performed on tissue samples. Neutralising antibody titres using quantitative SARS-CoV-2 virus neutralisation test (sVNT), pseudovirus and infectious virus neutralisation assays were performed on serum samples. SARS-CoV-2 serostatus was then correlated with the begging rank of each animal. Ex vivo, the ability of SARS-CoV-2 clinical isolates representative of ancestral and Omicron BA.1 variants to infect fallow deer precision cut lung slices and tracheal explants was evaluated.

### 2.2 Study area and population

The study population is located in the Phoenix Park, a 707 hectare urban park located in Dublin, Ireland, which receives an estimated 10 million visitors annually (Office of Public Works (OPW), official data) (*19*). There is a resident fallow deer population of approximately 600 free-ranging individuals (late summer 2018-2019 estimates inclusive of newborn fawns) (*19*). The behaviour of the animals and their interaction with humans has been extensively characterised with over 80% of the population individually identifiable with ear-tags (*21*). Of the total population, 86% of deer utilise areas of the park accessible to the public, whereas the remaining 14% (“avoiders’’) shun these human dense areas. Of the 86% of the deer that enter areas accessible to the public, 24% have a high contact rate with humans which includes taking food (“consistent beggars”), 68% display intermediate deer-human contact rates and food acceptance (“occasional beggars”), followed by deer systematically avoiding any interactions with humans despite living in areas open to the public (8%,“rare beggars”) (*19, 20, 22*). These categories are based on individuals’ begging rank, i.e. a scale running from most to least likely to beg based on previously-generated models of individual begging behaviour (Supplementary Information) (*19*).

This wild deer population lives in a large natural area surrounded by an encircling wall and the environs of a European capital city, and individuals occasionally disperse into surrounding residential areas adjacent to the park. While some animals have been killed in traffic collisions, they have no natural predators, with the only exception being red foxes who occasionally prey upon neonatal fawns (*21*). Deer are culled annually by professional stalkers over the winter period as part of the population management led by the Office of Public Works (OPW), the governmental body responsible for managing the deer in the park. The cull follows a defined culling strategy aimed at removing approximately 10% of the population whereby deer stalkers are required to target a number of animals based on sex and age to maintain a population that mimics a natural structure. Deer stalkers, however, are not instructed on which individuals (unique ear-tag codes) to remove, and culling operations are expected to randomly sample among the different begging categories and avoid artificial selection of behavioural traits (*19*). Ethical exemption for deer tagging and non-invasive behavioural observations was granted by University College Dublin’s Animal Research Ethics Committee (UCD AREC; approval number AREC-E-18-28-Ciuti).

### 2.3 Sample collection and storage

Culling of the fallow deer was carried out on the 2^nd^ and 25^th^ of November 2021, and on the 16^th^ of February 2022. In addition, serum samples collected following culling in November 2020 were obtained from archived samples (108 animals total; n=74 serum samples). Post-mortem, we sampled whole blood, retropharyngeal lymph nodes, palatine tonsil, nasopharyngeal mucosa and caecal content within one hour of death. The sampling strategy was chosen according to the highest viral loads reported from experimental infection of white-tailed deer (*12*). Ethical exemption for collection of post-mortem tissue was granted by UCD AREC (approval number AREC-E-21-49-Fletcher). Samples were transported to the laboratory immediately following collection, on ice, and tissue samples and caecal content stored at −80°C prior to RNA extraction. Whole blood samples were centrifuged at 1,500 x g, serum removed and frozen at −20°C prior to further analysis.

### 2.4 SARS-CoV-2 surrogate virus neutralisation test

A SARS-CoV-2 surrogate virus neutralisation test (sVNT) was performed on deer sera using the Genscript cPass^TM^ SARS-CoV-2 sVNT kit according to the manufacturer’s instructions. Briefly, sera were thawed and heat inactivated at 56°C for 30 minutes in a waterbath. Samples and controls were diluted 1:10 and mixed with horseradish peroxidase-receptor-binding domain (HRP-RBD) solution before incubating at 37°C for 30 minutes. Reaction mixtures were added to an ACE2-coated microtitre plate and incubated at 37°C for 15 minutes. The plate was washed and 3,3’,5,5’-tetramethylbenzidine (TMB) substrate was added to each well and incubated in the dark at 25°C for 15 minutes. The reactions were then quenched with stop solution before immediately reading the samples on a plate reader (Clariostar, BMG Labtech). Semi-quantitative results were expressed as percentage neutralisation. Sera were screened in duplicate within each assay, and two independent assays were performed to evaluate the replicability of results. The kit is not species specific and has been used as a screening tool to identify white-tailed deer and other mammalian species that are seropositive for SARS-CoV-2 (*12, 15*).

### 2.5 Nucleic acid extraction and SARS-CoV-2 qRT-PCR

Total RNA was extracted from tissues following homogenisation in TRIzol followed by extraction using the RNeasy Mini Kit (Qiagen, Germany) according to the manufacturer’s instructions. Briefly, a small amount of tissue (>25 mg) was added to 1 mL of TRIzol with a single 5mm stainless steel bead (Qiagen, Germany) and homogenised using a TissueLyser II (Qiagen, Germany) at maximum speed for 2 minutes. Following this, 0.1 mL of 1-bromo-3-chloropropane was added to each homogenised sample and shaken vigorously before incubating for 3 minutes. The sample was then spun in a pre-cooled centrifuge (4°C, 12,000 x g for 15 minutes) then 0.45 mL of the upper aqueous layer was transferred to a new tube and combined with 1 volume of 70% ethanol and mixed gently. The sample was added to RNeasy spin columns and the manufacturer’s protocol of the RNeasy Mini Kit was then followed. Samples were spiked with 1µg MS2 phage RNA as extraction efficiency controls.

An alternative protocol was used for extraction of RNA from caecal content samples which was adapted from Grierson et al (*23*). Briefly, 1-part caecal content was added to 9-parts 50 mM Tris-HCl (pH 8.0). This mixture was clarified by centrifugation (4°C, 4000 x g for 50 minutes) and the supernatant sequentially filtered through 0.45 µm and 0.22 µm filters. Filtrate was concentrated using 100 kDa Amicon (Merck-Millipore, Germany) spin columns. Free nucleic acids were digested using OmniCleave Endonuclease (Lucigen, USA) for 30 minutes. The same steps were then followed as with the tissue extraction protocol following the homogenisation step. As this protocol included a digestion step the same extraction control could not be used with the tissue extractions and samples were spiked with 100 µL of murine hepatitis virus (4.22 x 10^7^ TCID mL^-1^) as an extraction control.

qRT-PCR assays were performed on the ABI7500 Real-Time PCR System (Thermo Fisher Scientific, USA) in a total volume of 25 µL containing 5 µL of template employing Taqman Fast Virus 1-Step Master Mix (Thermo Fisher Scientific, USA). Oligonucleotide primer and probe sequences and thermocycling conditions are detailed in Table S2. A synthetic composition of ssRNA fragments of the SARS-CoV-2 genome (EURM-019, EC-JRC, Belgium) was used as a standard for SARS-CoV-2 E gene quantification. All samples and controls were analysed in triplicate with four triplicate quantification standards included in each qRT-PCR plate.

### 2.6 Cell culture, SARS-CoV-2 pseudovirus and infectious virus neutralisation test

VeroE6 cells (ATCC CRL-1586) and VeroE6 cells transiently expressing an untagged TMPRSS2 cDNA expression vector (Sinobiological, China; HG13070-UT) were propagated in Dulbecco’s Modified Eagle Medium (DMEM) supplemented with 10% fetal bovine serum (FBS), 2 mM L-glutamine (Gibco; Thermo Fisher, USA), and 1% non-essential amino acids (Gibco, Thermo Fisher, USA), as previously described (*24*). SARS-CoV-2 pseudoviruses (SARS-CoV-2pp) bearing spike proteins from variants of concern (VOC) Alpha, Delta, Omicron BA.1 and Omicron BA.2 were purchased from InvivoGen (Toulouse, France). SARS-CoV-2pp were generated as previously described, with some modifications (*25*). Briefly, pseudoviruses were generated by co-transfecting 293T cells with plasmids encoding a HIV-1 provirus expressing luciferase (pNL4-3-Luc-R-E-) and plasmids containing vesicular stomatitis virus glycoprotein (VSV-G), SARS-CoV-2 spike protein or a no envelope control (Env-). Supernatants were harvested 48 h and 72 h post transfection, filtered through a 0.45 µm filter and stored at −80°C.

SARS-CoV-2 (2019-nCoV/Italy-INMI1 from EVA Global; GenBank accession MT077125.1) was propagated in VeroE6 cells. All virus used in this study was at passage 3. Cells were inoculated with an MOI 0.01 for 2 hours, washed with phosphate-buffered saline (PBS), and the medium replaced with DMEM containing 2% FBS. When cultures were fully infected [cytopathic effect (CPE) observed in >80% of cells], flasks were freeze-thawed three times and supernatants collected and clarified at 3500 rpm for 30 minutes at 4°C. Supernatant was collected, aliquoted and stored at −80°C. TCID_50_ assays were performed on VeroE6 cells in quadruplicate and infectious titre determined using the Reed-Muench method. A SARS-CoV-2 Omicron BA.1 (CEPHR_IE_BA.1_0212, GenBank accession ON350968, Passage 2) clinical isolate, isolated from a SARS-CoV-2 positive nasopharyngeal swab from the All-Ireland

Infectious Disease (AIID) cohort (*26*), was isolated and amplified on Vero E6/TMPRSS2 cells (#100978), obtained from the Centre For AIDS Reagents (CFAR) at the National Institute for Biological Standards and Control (NIBSC) (*27*).

Pseudovirus or infectious virus was incubated 1:1 with deer sera at 37°C for 1 hour, and then titrated on VeroE6/TMPRSS2 cells. After 48 hours, for pseudovirus infections, cells were lysed with passive lysis buffer (Promega) and luciferase activity quantified using a luminometer (TriStar^2^ LB 942 Multimode Reader; Berthold Technologies, Germany). Infectivity was expressed as relative light units minus the signal for the no envelope control, and infectivity expressed relative to the virus only control. For infectious virus assays, CPE was scored 48 h post-infection and TCID_50_ calculated according to the method of Reed and Muench (*28*). To differentiate between CPE and cytotoxicity following incubation with sera, each serum sample was titrated alone on VeroE6/TMPRSS2 cells at the same dilutions as used for TCID_50_ assays. Cells were scored for cytotoxicity 48 h post-inoculation, and only dilutions that demonstrated an absence of visible cytotoxic effect (i.e. rounded or detached cells) were scored in TCID_50_ assays.

### 2.7 Determination of SARS-CoV-2 superlineages circulating during the sampling months

SARS-CoV-2 nanopore whole-genome sequencing (WGS) from human clinical samples collected in the Republic of Ireland was performed as described previously (*29*), covering periods of one month contemporaneous with the deer culling dates were downloaded from GISAID (https://gisaid.org/) and the PANGOLIN lineage was assigned using the PANGOLIN application v4.3 and pangolin data v1.20. The superlineages were assigned using the most recent common ancestors of each of the considered sequences with B.1, B.1.177, B.1.1.7 (Alpha), P.2 (Zeta), B.1.617.2 (Delta), B.1.1.529.1 (Omicron BA.1), B.1.1.529.2 (Omicron BA.2) and B.1.1.529.3 (Omicron BA.3). For graphical representations, a subset with a total of 108 genomes were sampled proportional to their superlineage frequencies from the three sampling periods.

The SARS-CoV-2 genome sequences were multiple-sequence aligned with the reference sequence (GenBank: MN908947) by the MAFFT program using the FFT-NS-I algorithm (*30*). A phylogenetic tree was inferred with the aligned sequences (n=109) by the program RAxML with a general time reversible (GTR) substitution model and the support for the branches estimated with a bootstrap approach with 100 repetitions.

### 2.8 Generation and SARS-CoV-2 infection of fallow deer tracheal explants and precision cut lung slices

Fallow deer lungs from two seronegative animals were removed immediately following culling, one lung clamped across the mainstem bronchus using a haemostat and transported to the laboratory on ice. Each lung was perfused with 2% low melting temperature agarose (Sigma-Aldrich, US) via the mainstem bronchus and allowed to set for approximately 10 minutes at room temperature. A 2 cm x 2 cm x 1 cm (LxWxH) section of agarose infused lung was dissected and embedded in an agarose mould and cut into 250 µm slices using a Leica VT1000S Vibratome (Leica Biosystems, Germany). Precision Cut Lung Slices (PCLS) of identical diameter were then generated using an 8 mm biopsy punch. Slices were transferred to individual wells of a 24 well plate and cultured in 2 mL Lung Slice Media (LSM), as described by Cousens et al. (*31*). Tracheal tissue from two seronegative animals was removed from the upper third of the trachea at culling, transported to the laboratory on ice, resected and cut longitudinally, through the trachealis muscle. The mucosa was carefully dissected away from the cartilage and an 8 mm biopsy punch was used to generate tracheal explants. Tracheal explants were cultured on the apical surface of Corning Transwell polycarbonate inserts with 0.4 μm pores (Catalogue 10482181) with 2 mL culture media added to the basolateral side of the models, creating an air-liquid interface. Tracheal explants were cultured in Tracheal Explant Media (TEM) consisting of LHC media supplemented with 5% heat-inactivated foetal calf serum, 50 U/mL penicillin/streptomycin, 1.25 μg/mL amphotericin B, and 1x Insulin-Transferrin-Selenium solution (Gibco).

Fallow deer PCLS and tracheal explants from two deer were inoculated with SARS-CoV-2 Italy-INMI-1, representing the ancestral virus, or an Omicron BA.1 clinical isolate as described in Section 2.5. For tracheal explant infections, 100 μL virus (titres: Italy-INMI-1: 3.9 x 10^5^ TCID mL^-1^; Omicron: 1.3 x 10^3^ TCID mL^-1^) was added to the apical (epithelial) surface of the tracheal tissue, and for PCLS, 500 μL virus was added to each well containing 500 μL culture media. 24 h post-inoculation, virus was removed and tracheal and PCLS tissue was washed 5x with warm PBS. Culture media was replaced and models were incubated for 72 h, after which tissue was fixed in 10% neutral buffered formalin for 72 h.

### 2.9 Immunohistochemistry to detect SARS-CoV-2 antigen in deer tissue

Formalin-fixed, paraffin-embedded (FFPE) fallow deer trachea and lung sections were immunohistochemically stained to evaluate the expression profile of SARS-CoV-2. Consecutive sections were cut at 4 µm thickness. A mouse monoclonal IgG1 antibody (1A9; GTX632604; GeneTex, USA) was used as the primary antibody. Staining was performed using the DAKO Link 48 Autostainer, as per the manufacturer’s instructions (Agilent Technologies Inc., USA) and the EnVision Flex kit (K8002, DAKO, Agilent Technologies Inc., US). Antigen retrieval was performed using DAKO PTLink (Agilent Technologies Inc., USA) for 20 min at 97°C in target retrieval buffer, pH 6 (K8005, DAKO, Agilent Technologies Inc., USA). Tissue sections were blocked for endogenous peroxidase and non-specific binding by incubating with 30% H_2_O_2_ (H1009, Sigma Aldrich, USA) for 30 min and protein block (X0909, DAKO, Agilent Technologies Inc., US) and T20 buffer (#37543, Thermo Fisher Scientific, USA) for 15 mins each. Optimal dilution of SARS-CoV-2 antibody was determined, using a FFPE, cell pellet of VeroE6 cells infected with SARS-CoV-2 Italy-INMI-1, and uninfected cell pellet as positive and negative controls, respectively. The optimal dilution and incubation time was 1:1000 for 30 min. Tissue sections were then washed and incubated with HRP for 20 min. The chromogen 3,39-diaminobenzidine (DAB) was used for revelation (twice for 5 min). Negative controls were run under identical conditions for each case, with diluent in place of primary antibody. An isotype control (IHC universal negative control reagent ADI-950-231-0025, Enzo Life Sciences, UK) was also included for each sample. Slides were counterstained with haematoxylin (K8008, DAKO, Agilent Technologies Inc., USA), and rinsed in deionized water. Slides were dehydrated, permanently mounted, scanned and digitised using the Aperio AT2 Digital Slide Scanner (Leica Biosystems, Germany), and images were reviewed using Aperio ImageScope 12.4 software (Leica Biosystems, Germany).

### 2.10 Statistical Analyses

*In vitro* results are expressed as the mean ±1 standard deviation of the mean (SD), except where otherwise stated. Statistical analyses were performed using one-way ANOVA in Prism 9.0 (GraphPad).

## 3. Results

### 3.1 Fallow deer are seropositive for SARS-CoV-2

Using a surrogate sVNT specific to SARS-CoV-2 (Genscript cPass SARS-CoV-2 sVNT), we screened fallow deer sera from animals culled in November 2020, November 2021 and February 2022 for neutralising antibodies to SARS-CoV-2. All animals culled in November 2020 were below the cut-off of the assay (30% neutralisation) and so were considered seronegative. All animals, with the exception of one, from the November 2021 cull were also seronegative. Animal 18_B_2021, a 4 year old male culled on 25^th^ November 2021, had a neutralisation value of 30% in the sVNT assay so is considered seropositive. In addition, 12/21 (57%) animals culled in February 2022 were seropositive for SARS-CoV-2, with percent neutralisation values >30% (Fig 1; Table 1). SARS-CoV-2 seropositive animals were a range of ages ranging from a fawn (<1 year) to mature animals eight years of age, and from a variety of sub-herds within the park (Table 1). Of the 13 seropositive animals, 10 were male (77%; Table 1).

**Figure 1.**
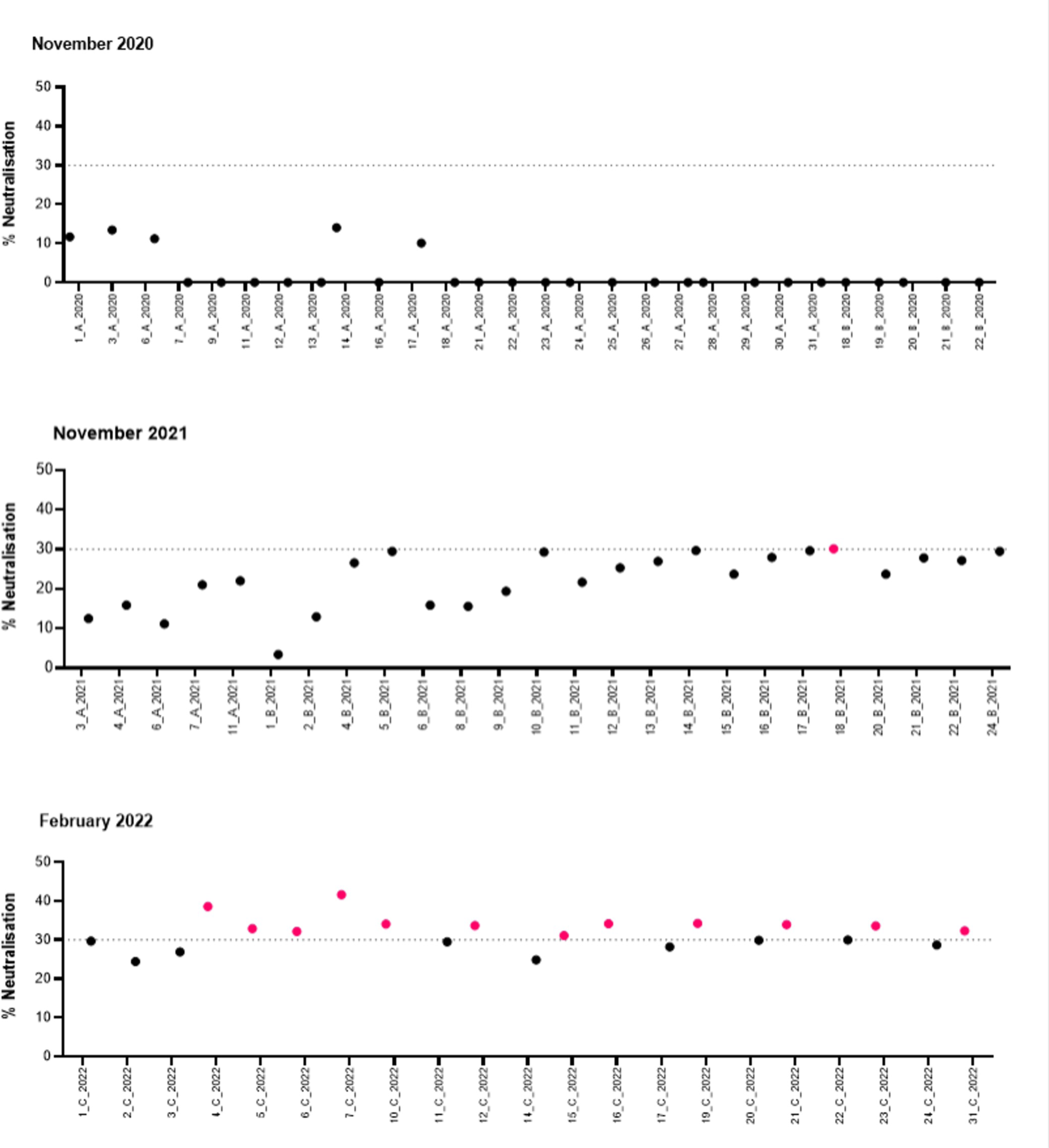
Fallow deer are seropositive for SARS-CoV-2. Serum from fallow deer was screened in duplicate for SARS-CoV-2 neutralising antibodies using the Genscript cPass SARS-CoV-2 sVNT assay. The dashed lines indicate a cut-off of 30% neutralisation, at or above which sera are considered seropositive for SARS-CoV-2. Data is presented as mean percentage neutralisation from duplicate wells from two independent assays.

**Table 1.** Profile and SARS-CoV-2 PCR status of seropositive fallow deer. SARS-CoV-2 seropositive fallow deer sampled in February 2022 were PCR negative for SARS-CoV-2 E (envelope) gene on all tissues tested (retropharyngeal lymph nodes, nasopharyngeal mucosa, palatine tonsil and caecal content). Age is presented in years (y). Begging ranks (occasional or consistent beggars, or unranked indicating that the animals are untagged) are presented for each animal. ND: not detected.

| Animal Code | Sample Date | Sex | Age (y) | SARS-CoV-2 sVNT (% neutralisation) | Tissue PCR | Caecal Content PCR | Begging Category |
| --- | --- | --- | --- | --- | --- | --- | --- |
| 18_B_2021 | 25/11/2021 | M | 4 | 30 | Undetected | Undetected | Unranked |
| 4_C_2022 | 16/02/2022 | M | 1 | 39 | Undetected | Undetected | Occasional |
| 5_C_2022 | 16/02/2022 | F | 6 | 33 | Undetected | Undetected | Occasional |
| 6_C_2022 | 16/02/2022 | M | 0 | 32 | Undetected | Undetected | Unranked |
| 7_C_2022 | 16/02/2022 | M | 1 | 42 | Undetected | Undetected | Occasional |
| 10_C_2022 | 16/02/2022 | F | 8 | 34 | Undetected | Undetected | Occasional |
| 12_C_2022 | 16/02/2022 | M | 2 | 34 | Undetected | Undetected | Unranked |
| 15_C_2022 | 16/02/2022 | F | 3 | 31 | Undetected | Undetected | Occasional |
| 16_C_2022 | 16/02/2022 | M | 3 | 34 | Undetected | Undetected | Consistent |
| 19_C_2022 | 16/02/2022 | M | 7 | 34 | Undetected | Undetected | Unranked |
| 21_C_2022 | 16/02/2022 | M | 5 | 34 | Undetected | Undetected | Consistent |
| 23_C_2022 | 16/02/2022 | M | 5 | 34 | Undetected | Undetected | Unranked |
| 31_C_2022 | 16/02/2022 | M | 1 | 32 | Undetected | Undetected | Unranked |

### 3.2 Fallow deer are qRT-PCR negative for SARS-CoV-2

Real time qRT-PCR was used to quantify SARS-CoV-2 E gene in retropharyngeal lymph nodes, palatine tonsil, nasopharyngeal mucosa and caecal content collected in November 2021 and February 2022. All animals were PCR negative for SARS-CoV-2, regardless of serostatus (Table 1 and Table S1). Positive ssRNA SARS-CoV-2 controls amplified in all assays, demonstrating that the qRT-PCR assay amplified SARS-CoV-2 E gene correctly (data not shown).

### 3.3 Begging behaviour of SARS-CoV-2 seropositive fallow deer

Notably, the majority of animals sampled at all timepoints were consistent or occasional beggars, taking food from humans (Fig 2A) (*19*). Of the SARS-CoV-2 seropositive animals sampled in February 2022, 7/12 (58%) were consistent or occasional beggars, with the remaining six animals unranked, indicating that they were not tagged and were therefore not individually identifiable or, in the case of 6_C_2022, was a tagged fawn with no assigned rank. However, this fawn’s mother (Yellow663) is a documented consistent beggar (*19*), so it is possible that this fawn came in contact with humans while following its mother. Beggars took a variety of foodstuffs from humans (Fig 2B) and previous work has demonstrated that park visitors pet the deer, particularly around the nose and face (*19*). Petting of males is more common than females as males are bolder and tend to tolerate greater levels of contact and proximity with humans (*19*). As the culling process was random, the subset of culled individuals that we sampled were representative of the entire population in terms of begging rank (Fig 2C), begging category as an alternative metric to begging rank (Fig 2D) (*19*), and age and sex classes (Fig 2E). We opted to use begging categories, rather than begging rank, as our basis for comparison as begging categories are computed using the errors generated by the model, as opposed to begging ranks (i.e. the random intercepts produced by the model used to calculate begging behaviour for each individual) (Supplementary Information) which do not account for these errors (*19*).

**Figure 2.**
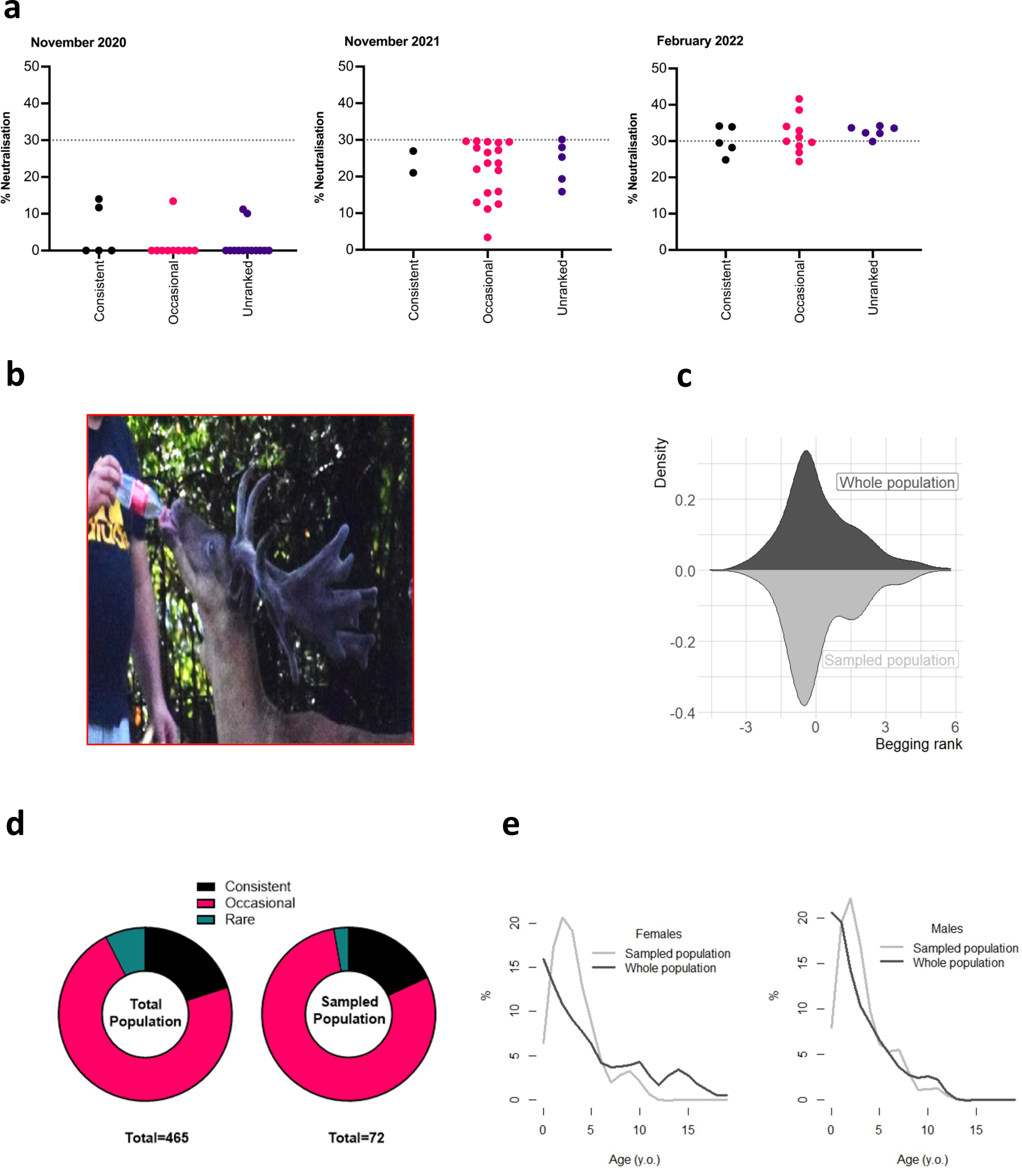
Begging behaviour of Fallow deer sampled for SARS-CoV-2 neutralizing antibodies. A) Seropositive fallow deer sera % neutralisation (y axis) plotted against begging rank (x axis) for each serum sample analysed. B) Example of fallow deer-human interaction. C) Begging rank distributions in the whole deer population in comparison to the sampled individuals. D) Proportion of whole population compared to the sampled population split by begging category. E) Age and sex structure of the entire population compared with the sampled individuals.

### 3.4 SARS-CoV-2 superlineages circulating in the human population during the deer sampling months

The SARS-CoV-2 genome sequences (n=5012) from human clinical samples circulating during the sampling months November 2020 (n=224), November 2021 (n=2883) and February 2022 (n=1905) allowed us to contextualise the different waves of variants circulating in the human population at the time of the deer culls (Fig 3). In addition, these genome sequences demonstrated the clear demarcation between the different waves of variants and non-overlapping phylogenetic relationships among the lineages identified in the three sampling periods. During the first sampling date (November 2020), B.1 and B.1.177 were the major variants circulating amongst the Irish population, with variants Alpha and Zeta detected during this month, and the former predominating in the months afterwards (*26*). In November 2021, the Delta variant was the major circulating variant for the entire month, whereas in February 2022, Omicron variants BA.1 and BA.2 were the major variants detected, with BA.3 detected at the end of the month.

**Figure 3.**
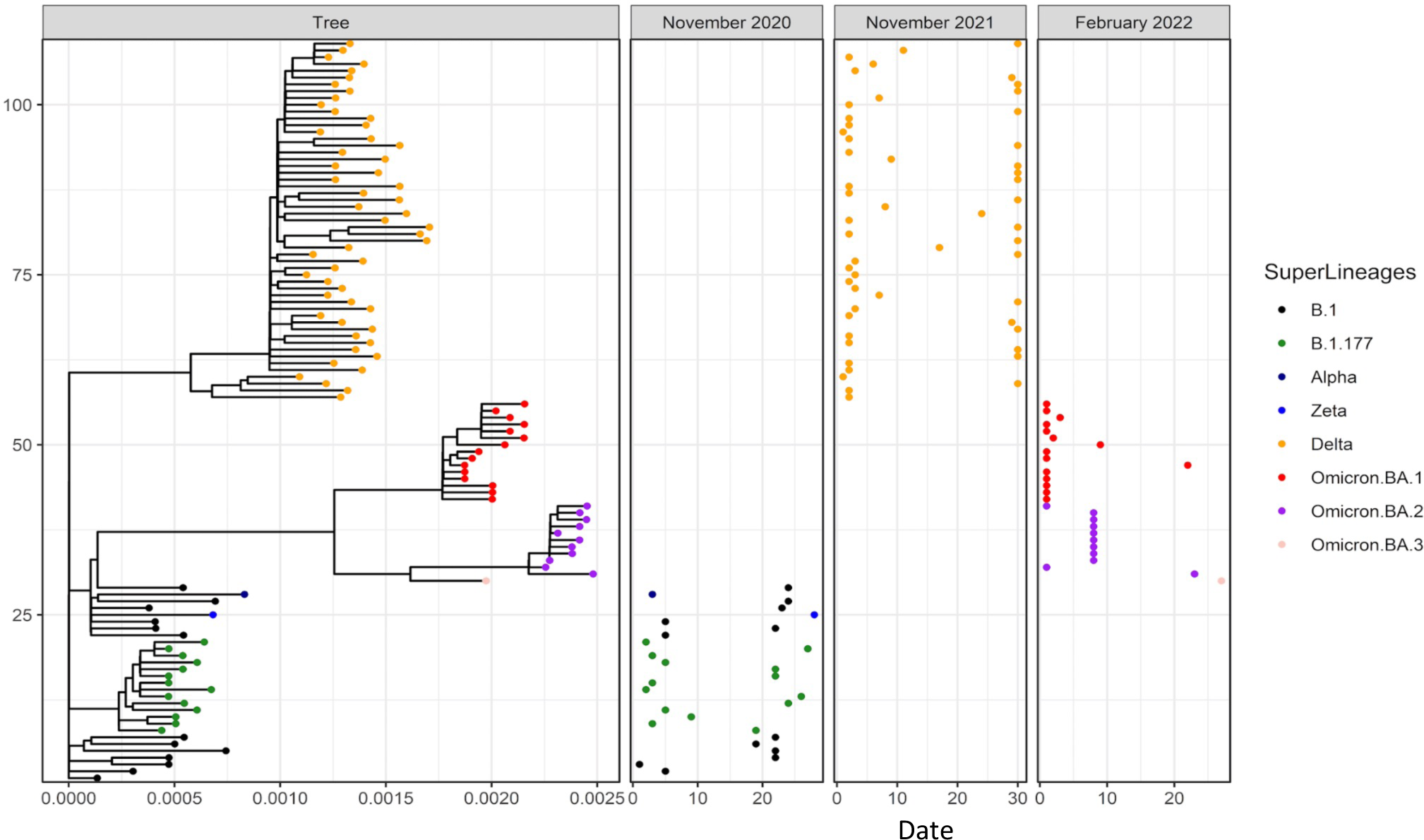
SARS-CoV-2 superlineages circulating during the sampling months. SARS-CoV-2 whole-genome sequences from human clinical samples collected in the Republic of Ireland covering months corresponding to the deer culling dates (November 2020, November 2021 and February 2022). PANGOLIN lineages are shown with corresponding major circulating variants for each month. The branch lengths of the phylogenetic tree show the number of base substitutions per site as shown in the scale at the bottom of the tree. The location of dots shown in each month corresponds to the sampling date in each month (horizontal axis) and the phylogenetic position in the tree panel (vertical axis).

### 3.5 SARS-CoV-2 sVNT positive sera neutralise SARS-CoV-2 pseudovirus and infectious virus

To confirm the ability of sVNT positive fallow deer sera to neutralise SARS-CoV-2pp and infectious virus, four seropositive sera with the highest neutralisation titres from the sVNT from the February 2022 cull were selected, together with a serum sample from the November 2021 cull that was at the cut-off to be considered seropositive (18_B_2021). All seropositive sera neutralised pseudoviruses bearing Alpha, Delta, Omicron BA.1 and BA.2 spike proteins, whereas 18_B_2021 only significantly inhibited BA.2 (Fig 4A). None of the five sera neutralised VSV-Gpp, an unrelated pseudovirus (data not shown). Using an infectious virus isolate representing the ancestral SARS-CoV-2 virus (Italy-INMI1) and an isolate of Omicron BA.1, significant inhibition of the ancestral virus by three of five fallow deer sera, and by all five sera in the case of Omicron BA.1 (Fig 4B; Table S3).

**Figure 4.**
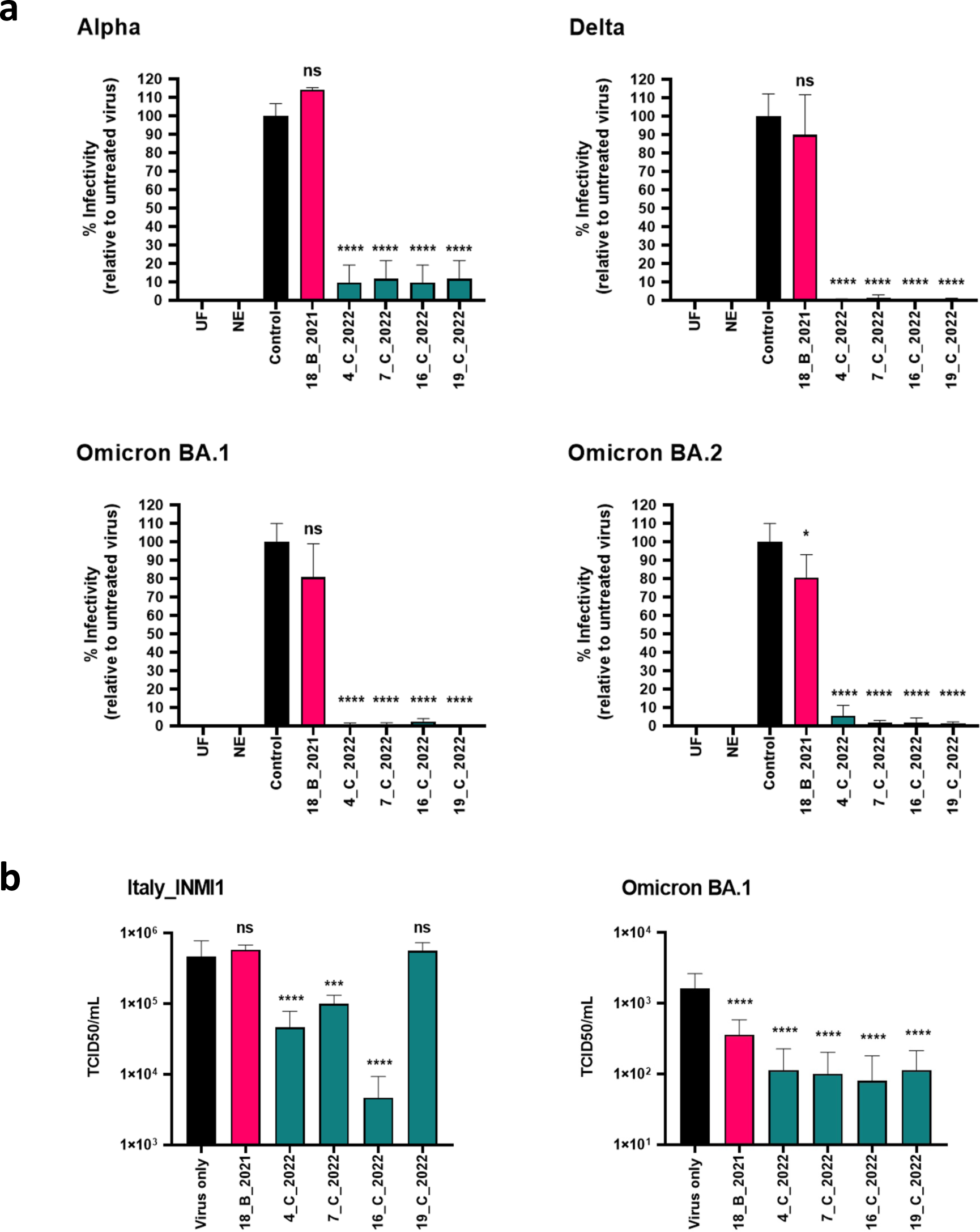
cPass positive deer sera neutralize SARS-CoV-2 VOC. A) SARS-CoV-2 pseudoviruses were incubated with seropositive deer sera in a 1:1 ratio in triplicate and then used to infect VeroE6/TMPRSS2 cells. Control is virus incubated in a 1:1 ratio with culture media, in triplicate. Relative light units (RLU) from each condition are expressed relative to the control, untreated virus as % infectivity. UF: uninfected cells; NE: no envelope naked pseudovirus control. Data are representative of two independent experiments, n=3 biological replicates per experiment. B) Deer sera were incubated with the infectious SARS-CoV-2 isolate Italy_INMI1, representing the ancestral virus, and Omicron BA.1 in the same way as described in A. Data are represented as 50% tissue culture infectious dose (TCID_50_), calculated according to the method of Reed and Muench, 193828. Data are representative of two independent experiments, n=8 biological replicates per experiment (Supplementary Table 2). One-way ANOVA: ns=p>0.05, ***p<0.001, ****p<0.0001.

### 3.6 SARS-CoV-2 infects fallow deer trachea and lung ex vivo

Two board certified pathologists (AF; medical pathologist and JC; veterinary pathologist) reviewed the digital slides and assessed the presence or absence of SARS-CoV-2 staining and the likely cells demonstrating immunoreactivity. Both pathologists evaluated the slides without prior knowledge of sample treatment. In tracheal explants challenged with SARS-CoV-2 Italy-INMI1 (isolate representative of ancestral SARS-CoV-2), occasional antigen positive cells within tracheal epithelium was observed in one animal (Fig 5A), whereas no immunoreactivity was detected in tracheal tissue challenged with Omicron BA.1. In contrast, cells morphologically consistent with type 2 pneumocytes were consistently observed to be immunoreactive in lung tissue from both animals inoculated with Omicron BA.1, but not with Italy-INMI1 (Fig 5B). No immunoreactivity was observed in tracheal or lung tissue from IgG controls (Fig 5B) or mock-challenged tissue (data not shown).

**Figure 5.**
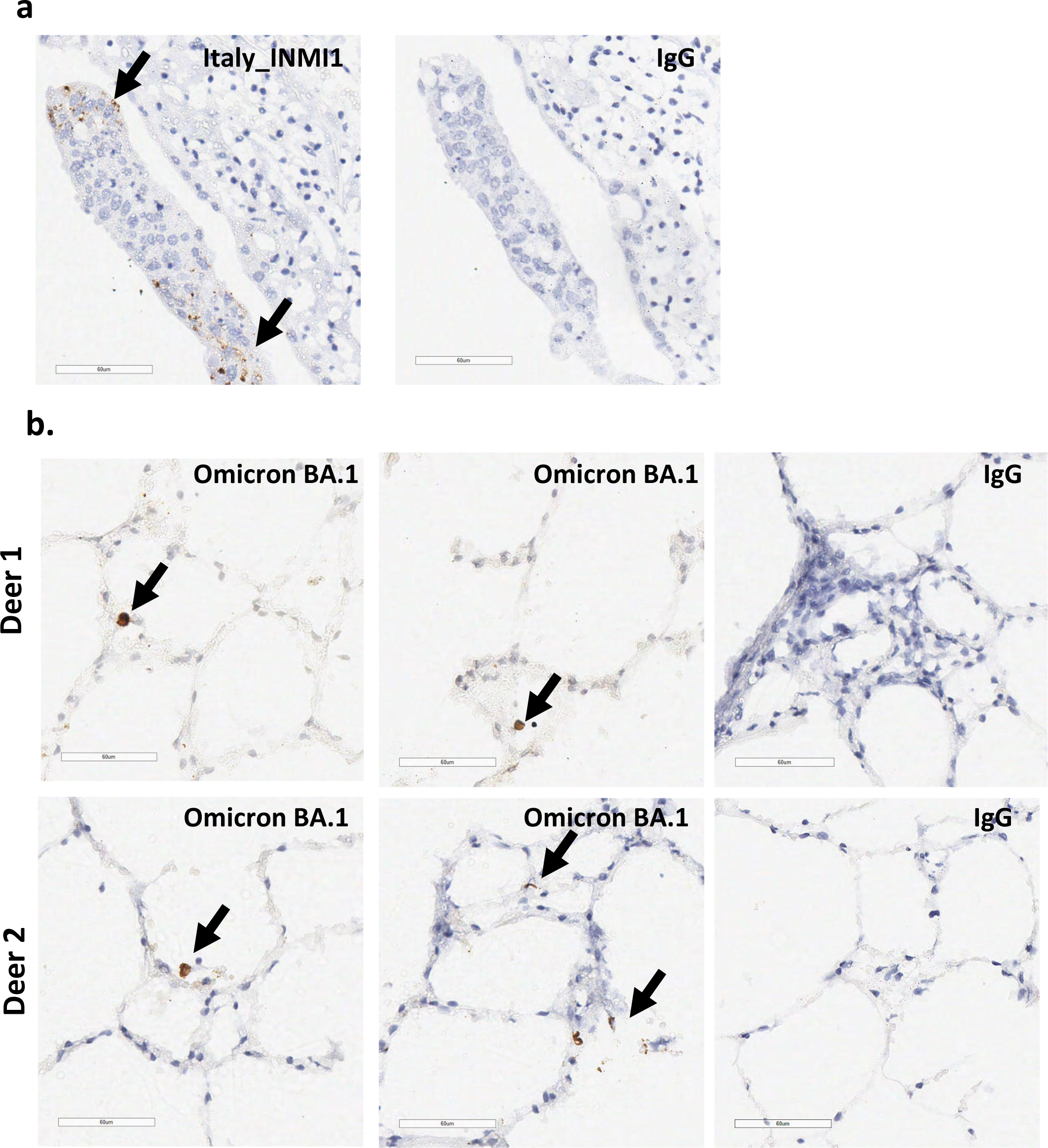
SARS-CoV-2 infects fallow deer trachea and precision cut lung slices *ex vivo.* **A)** Tracheal explant (deer 1) inoculated with SARS-CoV-2 Italy-INMI1, demonstrating viral antigen immunoreactivity in tracheal epithelium (arrows; left image). No immunoreactivity was observed in the IgG control (right image). B) Precision cut lung slices (PCLS) from deer 1 and 2 inoculated with SARS-CoV-2 Omicron BA.1, demonstrating immunoreactivity in cells morphologically consistent with type 2 pneumocytes (arrows). No immunoreactivity was observed in the IgG control for either animal.

## 4. Discussion

The World Health Organisation (WHO), World Organisation for Animal Health (WOAH) and Food and Agriculture Organisation (FAO) have emphasised the importance of continued monitoring of SARS-CoV-2 in wildlife, given the potential importance of wildlife species in the establishment and propagation of animal reservoirs and generation of novel variants with altered transmissibility and/or pathogenicity (*32*). The importance of minimising the risk of transmission of SARS-CoV-2 between humans and wildlife has also been emphasised, including educating the public about contact with wild animals and the risks associated with feeding these wild species (*32*).

In our study, interactions with the fallow deer population in the Phoenix Park have increased in the last ten years, and this has been attributed to increased social media visibility of the animals and their willingness to take food from humans (*19*). During the COVID-19 pandemic, there were initial increases in human-deer interactions in 2020, due to increased numbers of people using the park for recreation during lockdown, but this decreased to pre-pandemic levels by summer 2021 (*33*). To date, white-tailed deer (*Odocoileus virginianus*) are the only deer species that have been reported to be susceptible to SARS-CoV-2 infection. Multiple studies have reported high levels of PCR-positive white-tailed deer, with up to 30-40% of free-ranging animals reported to be PCR positive (*9*). Importantly, white-tailed deer are capable of shedding infectious virus that can transmit between deer (*12*) and deer-to-human transmission has been documented (*34, 35*). Other mammalian species that can transmit SARS-CoV-2 include mink, raccoon dogs, cats, ferrets, hamsters, mice, Egyptian fruit bats and deer mice(*4*).

In the present study, we initially screened sera from fallow deer for SARS-CoV-2 neutralising antibodies using a commercial sVNT assay (Genscript cPass). Of the 13 seropositive animals, the highest percent neutralisation was 42%, with a mean of 34% neutralisation in our fallow deer population. Using the same sVNT assay, reported neutralisation levels for white-tailed deer approach 80-90% (*7*), and similar levels have been reported for animal species susceptible to SARS-CoV-2 infection, including hamsters, mink, ferrets and cats, and humans (*36*). In the present study, SARS-CoV-2 neutralisation levels using pseudovirus was higher than that observed using the sVNT assay in the present study for sera from February 2022. The positive serum from November 2021 (18_B_2021), which had a sVNT neutralisation titre at the cut-off of 30%, significantly inhibited BA.2 pseudovirus but not pseudoviruses bearing spike proteins from other variants. Moreover, 18_B_2021 significantly inhibited infectious Omicron BA.1 but not the ancestral virus. The sera from the February 2022 cull had higher titres in the sVNT assay than that of the November 2021 serum sample, and this was reflected in the ability of these sera to neutralise pseudoviruses. However, differences in neutralisation titres for the ancestral virus and Omicron BA.1 were observed for several seropositive sera, with all sera capable of neutralising Omicron BA.1 but only three of five significantly neutralising Italy_INMI1. The sVNT uses SARS-CoV-2 spike protein from the ancestral Wuhan virus. It is possible that we would have observed higher neutralisation levels using the sVNT test if spike proteins from other SARS-CoV-2 variants had been incorporated. This result is in agreement with other studies, highlighting that the sVNT assay should be interpreted qualitatively, rather than quantitatively and that differences in assays evaluating SARS-CoV-2 neutralising antibodies have been reported previously (*36*). The sVNT assay specifically detects SARS-CoV-2 neutralising antibodies, with no cross-reactivity to other human coronaviruses or respiratory viruses, or to MERS-CoV (*36*) with the exception of SARS-CoV, which is closely related antigenically to SARS-CoV-2 (*37*). However, during the timeframe the fallow deer were sampled in the current study, during the SARS-CoV-2 pandemic when there was no evidence of SARS-CoV circulating in Ireland, the likelihood of these animals harbouring neutralising antibodies to SARS-CoV is considered negligible.

While SARS-CoV-2 seropositive animals in the present study were ranked as mainly occasional beggars, with fewer animals frequent beggars or unranked (*19*), this distribution is representative of the deer population as a whole. In total, consistent beggars represent 24% of the observed population, while occasional beggars represent 68% of the population and rare represent 8%. Overall, 86% of the population’s begging rank is categorised (*19*). The remaining subset of the population (14%, avoiders) limit themselves to smaller areas of the park that are closed to the public. We only sampled two rare beggars or avoiders in the present study, likely due to their avoidance of contact with humans and therefore their lower likelihood of being culled. This is in agreement with reports from random capture studies, i.e., that we are less likely to trap, or in this case observe, the shy-inactive individuals present in any population due to their careful avoidance of humans (*19, 38-41*). Overall, 77% of the seropositive animals in this study were male, and in total, 60% of the animals sampled in February 2022 were male. Previous studies have demonstrated that males in this population are more likely to beg for food from humans (*19*), which indicates that begging behaviour was positively correlated with serostatus in this study consistent with the hypothesis of anthroponosis.

During the three sampling periods in the current study, there were diverse major viral variants circulating within the human population within Ireland as revealed by SARS-CoV-2 sequencing data generated over the course of the COVID-19 pandemic (*29, 42, 43*). Given the high frequency of interactions with humans, it is likely that the deer population was exposed to SARS-CoV-2 through contact with humans. While the February 2022 sampling period coincided with Omicron circulation in the human population, it is not possible to definitively state which subvariant the seropositive deer in this study were exposed to, or indeed whether there was transmission of SARS-CoV-2 between deer. The majority of surveillance studies investigating the serostatus of deer in Europe have not sampled animals post-2021 and have not studied deer with defined and quantified interactions with humans(*15, 16*). The present study, together with a recent report demonstrating ACE2 receptor expression in the respiratory tract of several mammalian and deer species, including fallow deer (*17*) highlights the need for ongoing surveillance particularly as novel VOCs are still emerging (*32*).

We demonstrated that fallow deer PCLS can support SARS-CoV-2 infection with a clinical isolate of Omicron BA.1, but not with Italy-INMI1, a SARS-CoV-2 isolate representing the ancestral virus. SARS-CoV-2-positive cells were observed in type 2 pneumocytes within PCLS 96 h post-inoculation in Omicron-challenged tissues. In contrast, occasional SARS-CoV-2-positive cells were observed in tracheal epithelium from one animal challenged with Italy-INMI-1. The tissue distribution of viral antigen-positive cells correlates with previously-reported ACE2 expression in the lungs of fallow deer (*17*), suggesting that Omicron may have increased propensity to utilise ACE2 as a receptor for SARS-CoV-2 entry relative to the ancestral virus. The significance of the rarely observed viral antigen within tracheal epithelial cells from one of two animals challenged with Italy-INMI1 is unclear and warrants further investigation. While SARS-CoV-2 immunoreactivity in lung tissue inoculated with Omicron BA.1 mirrors ACE2 expression previously reported (*17*), whether other receptors, such as TMPRSS2, are also necessary for SARS-CoV-2 infection of fallow deer cells is currently unknown. SARS-CoV-2 in experimentally-infected white-tailed deer, challenged with a human B.1 SARS-CoV-2 isolate, infected epithelial cells in both upper respiratory tract and lung tissue (*10*). In human patients with COVID-19, alveolar type 2 pneumocytes are the main cell type that expresses ACE-2 and are also the main targets for SARS-CoV-2 infection in the human lung (*44-46*). Damage to these cells following viral infection can result in reduced production of surfactant, alveolar injury and impaired lung function, similar to SARS-CoV (*46-48*). None of the deer sampled in this study were PCR positive for SARS-CoV-2, but given the relatively short duration of viral replication in other species compared with the extended persistence of antibodies, we were more likely to detect seropositive animals compared with those actively shedding virus. Neutralising antibodies persist in naturally-infected white-tailed deer for over 13 months (*49*). However, it is currently unknown whether fallow deer can be productively infected with SARS-CoV-2 *in vivo*, shed virus or transmit infection to other fallow deer or other species including humans, or whether other variants, such as Delta, have the ability to infect fallow deer. Together, these data suggest that Omicron may have an elevated propensity to infect fallow deer lung in contrast to ancestral SARS-CoV-2, highlighting the importance of ongoing surveillance of deer species as new VOCs emerge.

In conclusion, this is the first report, to our knowledge, of SARS-CoV-2 seropositivity in European deer species, and the first report of seropositivity in any deer species other than white-tailed deer. While one animal culled in November 2021 was seropositive, 57% of fallow deer were seropositive for SARS-CoV-2 in February 2022 when Omicron was the dominant circulating variant in the local population, and fallow deer lung tissue could be infected with Omicron, but not ancestral SARS-CoV-2, *ex vivo*. This study highlights the importance of ongoing surveillance to identify novel animal and wildlife reservoirs of SARS-CoV-2 and other zoonotic pathogens. It also highlights the ecological risk factors to public health of interactions between human and animal species in peri-urban settings.

## Author Contributions

Conceptualisation: KP, AR, SC and NFF.

Methodology: KP, HB, RH, AR, LLG, JMcC, SOR, VG, MC, GG and NFF

Investigation: KP, HB, RH, AR, LLG, JMcC, SOR, VG, MC, GG PWM, VG and NFF.

Review of pathology slides and expert opinions: JC and AF.

Writing: KP, HB, RH, AR, LLG, GG, SC and NFF.

Review and editing: All authors.

## Acknowledgements

The authors wish to acknowledge the University College Dublin Veterinary Medicine Containment Level 3 Laboratory and its managers, Prof Stephen Gordon, Dr John Browne and Dr Bridget Hogg, for facilitating this study. The authors acknowledge Mr Marc Farrelly, Ms Tiffany Morey and Dr Christopher Evans for technical assistance, and the Office of Public Works, Ireland.

## Data Availability Statement

All data needed to evaluate the conclusions in the paper are present in the paper and/or the Supplementary Materials.

## References

1. P. V’Kovski, A. Kratzel, S. Steiner, H. Stalder, V. Thiel, Coronavirus biology and replication: implications for SARS-CoV-2. Nat Rev Microbiol 19, 155–170 (2021).

2. https://covid19.who.int/.

3. S. J. Cleary et al., Animal models of mechanisms of SARS-CoV-2 infection and COVID-19 pathology. Br J Pharmacol 177, 4851–4865 (2020).

4. https://www.ecdc.europa.eu/en/publications-data/sars-cov-2-animals-susceptibility-animal-species-risk-animal-and-public-health.

5. P. Jia, S. Dai, T. Wu, S. Yang, New Approaches to Anticipate the Risk of Reverse Zoonosis. Trends Ecol Evol 36, 580–590 (2021).

6. C. S. Bakshi, A. J. Centone, G. P. Wormser, SARS-CoV-2 is Emerging in White-Tailed Deer and Can Infect and Spread Among Deer Mice Experimentally: What About Deer Ticks? Am J Med 135, 1395–1396 (2022).

7. J. C. Chandler et al., SARS-CoV-2 exposure in wild white-tailed deer (Odocoileus virginianus). Proc Natl Acad Sci U S A 118, (2021).

8. G. Di Guardo, Susceptibility of white-tailed deer to SARS-CoV-2. Vet Rec 189, 408–409 (2021).

9. V. L. Hale et al., SARS-CoV-2 infection in free-ranging white-tailed deer. Nature 602, 481–486 (2022).

10. M. Martins et al., From Deer-to-Deer: SARS-CoV-2 is efficiently transmitted and presents broad tissue tropism and replication sites in white-tailed deer. PLoS Pathog 18, e1010197 (2022).

11. P. M. Palermo, J. Orbegozo, D. M. Watts, J. C. Morrill, SARS-CoV-2 Neutralizing Antibodies in White-Tailed Deer from Texas. Vector Borne Zoonotic Dis 22, 62–64 (2022).

12. M. V. Palmer et al., Susceptibility of white-tailed deer (Odocoileus virginianus) to SARS-CoV-2. J Virol 95, (2021).

13. C. M. Roundy et al., High Seroprevalence of SARS-CoV-2 in White-Tailed Deer (Odocoileus virginianus) at One of Three Captive Cervid Facilities in Texas. Microbiol Spectr 10, e0057622 (2022).

14. https://www.woah.org/en/oie-statement-on-monitoring-white-tailed-deer-for-sars-cov-2/.

15. M. Holding et al., Screening of wild deer populations for exposure to SARS-CoV-2 in the United Kingdom, 2020-2021. Transbound Emerg Dis 69, e3244-e3249 (2022).

16. A. Moreira-Soto et al., Serological Evidence That SARS-CoV-2 Has Not Emerged in Deer in Germany or Austria during the COVID-19 Pandemic. Microorganisms 10, (2022).

17. F. Z. X. Lean et al., Tissue distribution of angiotensin-converting enzyme 2 (ACE2) receptor in wild animals with a focus on artiodactyls, mustelids and phocids. One Health 16, 100492 (2023).

18. Y. Zhang et al., Cross-species tropism and antigenic landscapes of circulating SARS-CoV-2 variants. Cell Rep 38, 110558 (2022).

19. L. L. Griffin et al., Artificial selection in human-wildlife feeding interactions. Journal of Animal Ecology 91, 1892–1905 (2022).

20. L. L. Griffin et al., Reducing risky interactions: Identifying barriers to the successful management of human-wildlife conflict in an urban parkland. People Nat 4, 918–930 (2022).

21. F. C. P.W.G. Mallon, G. Gonzalez, W. Tinago, A.A. Garcia Leon, M. McCabe, E. de Barra, O. Yousif, J.S. Lambert, C.J. Walsh, J.G. Kenny, E. Feeney, M. Carr, P. Doran, P.D. Cotter on behalf of the All Ireland Infectious Diseases Cohort Study and the Irish Coronavirus Sequencing Consortium., Whole-genome sequencing of SARS-CoV-2 in the Republic of Ireland during waves 1 and 2 of the pandemic. MedRxiv, (2021).

22. https://www.who.int/news/item/07-03-2022-joint-statement-on-the-prioritization-of-monitoring-sars-cov-2-infection-in-wildlife-and-preventing-the-formation-of-animal-reservoirs.

23. L. L. Griffin, Nolan, G., Haigh, A., Condon, H., O’Hagan, H., McDonnell, P., Kane, A. and Ciuti, S., How can we tackle interruptions to human-wildlife feeding management? Adding media campaigns to the wildlife manager’s toolbox. People Nat in press, (2023).

24. B. Pickering et al., Publisher Correction: Divergent SARS-CoV-2 variant emerges in white-tailed deer with deer-to-human transmission. Nat Microbiol, (2022).

25. B. Pickering et al., Divergent SARS-CoV-2 variant emerges in white-tailed deer with deer-to-human transmission. Nat Microbiol 7, 2011–2024 (2022).

26. C. W. E. Embregts et al., Evaluation of a multi-species SARS-CoV-2 surrogate virus neutralization test. One Health 13, 100313 (2021).

27. https://www.hsa.gov.sg/docs/default-source/hprg-mdb/psar-covid-19-puo-tests/ab01_genscript-cpass-sars-cov-2-neutralization-antibody-detection-kitdaa6d27f8f5446e1ae7d7a1fedbc3e69.

28. P. A. Biro, N. J. Dingemanse, Sampling bias resulting from animal personality. Trends Ecol Evol 24, 66–67 (2009).

29. M. Leclerc, Zedrosser, A., & Pelletier, F., Harvesting as a potential selective pressure on behavioural traits. Journal of Applied Ecology 54, 1941–1945 (2017).

30. M. Leclerc, A. Zedrosser, J. E. Swenson, F. Pelletier, Hunters select for behavioral traits in a large carnivore. Sci Rep 9, 12371 (2019).

31. D. Sloan Wilson, A. B. Clark, K. Coleman, T. Dearstyne, Shyness and boldness in humans and other animals. Trends Ecol Evol 9, 442–446 (1994).

32. E. P. T. Alan M. Rice, Stephen Bridgett, Behnam Firoozi Nejad, Jennifer M. McKinley, The COVID-19 Genomics UK consortium, National SARS-CoV-2 Surveillance & Whole Genome Sequencing (WGS) Programme, Declan T. Bradley, Derek Fairley, Connor G. G. Bamford, Timofey Skvortsov, David A. Simpson, SARS-CoV-2 introductions to the island of Ireland. MedRxiv, (2023).

33. T. Funk et al., Characteristics of SARS-CoV-2 variants of concern B.1.1.7, B.1.351 or P.1: data from seven EU/EEA countries, weeks 38/2020 to 10/2021. Euro Surveill 26, (2021).

34. L. J. Reynolds et al., SARS-CoV-2 variant trends in Ireland: Wastewater-based epidemiology and clinical surveillance. Sci Total Environ 838, 155828 (2022).

35. K. K. Acheampong et al., Subcellular Detection of SARS-CoV-2 RNA in Human Tissue Reveals Distinct Localization in Alveolar Type 2 Pneumocytes and Alveolar Macrophages. mBio 13, e0375121 (2021).

36. B. Huang, Mucins produced by type II pneumocyte: culprits in SARS-CoV-2 pathogenesis. Cell Mol Immunol 18, 1823–1825 (2021).

37. A. Jain, COVID-19 and lung pathology. Indian J Pathol Microbiol 63, 171–172 (2020).

38. M. M. Aboudounya, R. J. Heads, COVID-19 and Toll-Like Receptor 4 (TLR4): SARS-CoV-2 May Bind and Activate TLR4 to Increase ACE2 Expression, Facilitating Entry and Causing Hyperinflammation. Mediators Inflamm 2021, 8874339 (2021).

39. K. C. Chow, C. H. Hsiao, T. Y. Lin, C. L. Chen, S. H. Chiou, Detection of severe acute respiratory syndrome-associated coronavirus in pneumocytes of the lung. Am J Clin Pathol 121, 574–580 (2004).

40. S. A. Hamer et al., Persistence of SARS-CoV-2 neutralizing antibodies longer than 13 months in naturally infected, captive white-tailed deer (Odocoileus virginianus), Texas. Emerg Microbes Infect 11, 2112–2115 (2022).

41. B. Amin et al., In utero accumulated steroids predict neonate anti-predator response in a wild mammal. Funct Ecol 35, 1255–1267 (2021).

42. L. L. Griffin, et al., Does artificial feeding impact neonate growth rates in a large free-ranging mammal? Roy Soc Open Sci 10, (2023).

43. S. S. Grierson, S. McGowan, C. Cook, F. Steinbach, B. Choudhury, Molecular and in vitro characterisation of hepatitis E virus from UK pigs. Virology 527, 116–121 (2019).

44. K. Purves et al., A novel antiviral formulation containing caprylic acid inhibits SARS-CoV-2 infection of a human bronchial epithelial cell model. J Gen Virol 104, (2023).

45. N. F. Fletcher et al., A novel antiviral formulation inhibits a range of enveloped viruses. J Gen Virol 101, 1090–1102 (2020).

46. S. Matsuyama et al., Enhanced isolation of SARS-CoV-2 by TMPRSS2-expressing cells. Proc Natl Acad Sci U S A 117, 7001–7003 (2020).

47. M. H. Reed LJ, A simple method of estimating fifty per cent endpoints. American Journal of Hygiene 27, 493–497 (1938).

48. K. Katoh, J. Rozewicki, K. D. Yamada, MAFFT online service: multiple sequence alignment, interactive sequence choice and visualization. Brief Bioinform 20, 1160–1166 (2019).

49. C. Cousens, L. Gibson, J. Finlayson, I. Pritchard, M. P. Dagleish, Prevalence of ovine pulmonary adenocarcinoma (Jaagsiekte) in a UK slaughterhouse sheep study. Vet Rec 176, 413 (2015).

