## Supplementary Methods for "First Eurasian cases of SARS-CoV-2 seropositivity in a free-ranging urban population of wild fallow deer"

**1. Begging ranks and categories**

*Step 1 - Modelling begging ranks:*

Data analyses were performed in R 4.0.5 (*1*). Feeding data was collected as previously described (*2*), and all individuals observed for less than 1080 minutes (i.e. 3 observation sessions) were dropped in order to remove potential noise from underrepresented individuals. Our final dataset was composed of 19,451 rows, encompassing data from feeding collections performed from the start of June to end of July (Friday-Sunday) from 2019-2021. Each row in our dataset corresponded with an observation of an identified individual, the code for the herd that they were present in, whether or not they engaged with an interaction (binomial; 1 = a line for each time they engaged in a separate feeding interaction over the course of that herd observation, 0 = they were documented as being present in the herd but never approached any available feeding interactions), information on the interaction (i.e. how many people were involved), and all relevant spatial and temporal information (outlined in (*2*)).

We fit a generalised linear mixed-effects model (GLMM) with binomial distribution of errors following *a priori* structure using *glmer* function in *lme4* (*3*), with begging as response variable and deer ID and herd ID (observation number) as crossed random intercepts. Our model *a priori* structure was built using predictors of interest that we selected prior to the beginning of the data collection. We included deer age, sex, herd size, the total number of people that attempted to interact with the herd during the observation session, the day of the week (categorical: Friday, Saturday, Sunday), the time of the day, the amount of time the observer spent monitoring the herd (duration, i.e. sampling effort), the month of observation (categorical: June, July) and year of study (categorical: 2019, 2020, 2021) as explanatory variables in our model. All numerical predictors were scaled to improve model convergence, and were included as both single and quadratic terms in the model to allow for non-linear patterns.

We also included three two-way interactions in the model: sex and age, sex of the individual and total amount of people that attempted to interact with the herd, and sex and herd size, all of which were included in both their linear and quadratic forms (resulting in six interactions in total). For details on variable selection rationale and *a priori* expectations, see (*2*). All predictors included in the model structure were not collinear (|*r_p_*| < 0.7) (*4*). We then extracted the random intercepts estimated by the GLMM for each individual ID (sometimes known as conditional modes, or best linear unbiased predictors, BLUPs; (*5*)), which were highly repeatable across individuals (*2, 6*). This constituted a ranking system, ranging from the deer that showed the lowest begging rate to the one that showed the highest rate after taking all model predictors into account.

*Step 2 – Extracting begging categories*

We extracted the begging rank (i.e. random intercept or BLUP value) and 95% confidence intervals for each random intercept value for each deer ID, and then summarised them into behavioural categories. These behavioural categories were identified depending on how the random effects and related confidence intervals were distributed around the median random effect (zero, i.e., the median begging behaviour of the population). Ultimately, begging behaviour exists on a continuum, but for clarification we subdivided these into three categories; those whose random effect’s 95% CIs remained greater than zero (“consistent” beggars), overlapped zero (“occasional” beggars), or was less than zero (“rare” beggars). This approach considers and categorises the BLUPs (random intercept) while connecting them with the associated error (confidence intervals) (sensu *7, 8*). These “begging categories” were then associated with the ID of each culled individual.

Table S1.

Demographics, SARS-CoV-2 serostatus, begging rank and category of fallow deer

(See separate Table S1.xlsx file)

Table S2.

**SARS-CoV-2 qRT-PCR primers, probes and cycling conditions**

| **Target Gene** | **Primer/Probe Sequence (5’ - 3’)** | **Cycling Conditions** | **Reference** |
| --- | --- | --- | --- |
| SARS-CoV-2 Envelope Gene | E_Sarbeco_F: ACAGGTACGTTAATAGTTAATAGCGT  E_Sarbeco_R: ATATTGCAGCAGTACGCACACA  E_Sarbeco_P1: FAM-ACACTAGCCATCCTTACTGCGCTTCG-BBQ | RT (50°C – 300s),  95°C – 20s,  45 cycles (95°C – 15s, 59°C – 60s) | Corman et al., (2020) (*9*) |

F: forward primer, R: reverse primer, P1: Probe.

Table S3.

Statistical analyses

(See separate Table S3.xlsx file)

**References**

1.	R Core Team, in *R Foundation for Statistical Computing, Vienna, Austria.* (2021).

2.	L. L. Griffin *et al.*, Artificial selection in human-wildlife feeding interactions. *J Anim Ecol* **91**, 1892-1905 (2022).

3.	D. Bates, M. Machler, B. M. Bolker, S. C. Walker, Fitting Linear Mixed-Effects Models Using lme4. *Journal of Statistical Software* **67**, 1-48 (2015).

4.	C. F. Dormann *et al.*, Collinearity: a review of methods to deal with it and a simulation study evaluating their performance. *Ecography* **36**, 27-46 (2013).

5.	G. K. Robinson, That BLUP is a good thing: the estimation of random effects. *Statistical science*, 15-32 (1991).

6.	S. N. M. A STOFFEL, H SCHIELZETH, S GOSLEE, , rptR: repeatability estimation and variance decomposition by generalized linear mixed-effects models. *Methods in Ecology and Evolution* **8**, 1639-1644 (2017).

7.	J. D. Hadfield, A. J. Wilson, D. Garant, B. C. Sheldon, L. E. B. Kruuk, The misuse of BLUP in ecology and evolution. *The American Naturalist* **175**, 116-125 (2010).

8.	T. M. Houslay, A. J. Wilson, Avoiding the misuse of BLUP in behavioural ecology. *Behavioral Ecology* **28**, 948-952 (2017).

9.	V. M. Corman *et al.*, Detection of 2019 novel coronavirus (2019-nCoV) by real-time RT-PCR. *Euro Surveill* **25**, (2020).
